# Aging and MPTP-sensitivity depend on molecular and ultrastructural signatures of astroglia and microglia in mice substantia nigra

**DOI:** 10.1101/2020.12.15.422212

**Authors:** PL Abhilash, Upasna Bharti, Santhosh Kumar Rashmi, Mariamma Philip, T. R. Raju, Bindu M. Kutty, B.K. Chandrasekhar Sagar, Phalguni Anand Alladi

**Author notes:** **Corresponding Author:** Dr. Phalguni Anand Alladi, Senior Scientific Officer- Scientist ‘F‘, Department of Clinical Pharmacology and Toxicology. Formerly at- Department of Neurophysiology, National Institute of Mental Health and Neuro Sciences, Hosur Road Bangalore, India (Phalguni Anand Alladi).

## Abstract

Both astroglia and microglia show region-specific distribution in CNS and often maladapt to age-associated alterations within their niche. Studies on autopsied substantia nigra (SN) of Parkinson’s disease (PD) patients and experimental models propose gliosis as a trigger for neuronal loss. Epidemiological studies propose an ethnic bias in PD prevalence, since Caucasians are more susceptible than non-whites. Similarly, different mice strains are variably sensitive to MPTP. We had earlier likened divergent MPTP-sensitivity of C57BL/6J and CD-1 mice with differential susceptibility to PD, based on the numbers of SN neurons.

Here, we examined whether the variability was incumbent to inter-strain differences in glial features of male C57BL/6J and CD-1 mice. Stereological counts showed relatively more microglia and fewer astrocytes in the SN of normal C57BL/6J mice, suggesting persistence of an immune-vigilant state. MPTP-induced microgliosis and astrogliosis in both strains, suggests their involvement in pathogenesis. ELISA of pro-inflammatory cytokines in the ventral-midbrain revealed augmentation of TNF-α and IL-6 at middle-age in both strains that reduced at old-age, suggesting middle-age as a critical, inflamm-aging associated time-point. TNF-α levels were high in C57BL/6J, through aging and post-MPTP; while IL-6 and IL-1β were upregulated at old-age. CD-1 had higher levels of anti-inflammatory cytokine TGF-β. MPTP-challenge caused upregulation of enzymes MAO-A, MAO-B and iNOS in both strains. Post-MPTP enhancement in fractalkine and hemeoxygenase-1; may be neuron-associated compensatory signals. Ultrastructural observations of elongated astroglial/microglial mitochondria vis-à-vis the shrunken ones in neurons, suggest a scale-up of their functions with neurotoxic consequences. Thus, astroglia and microglia modulate aging and PD-susceptibility.

**Highlights:**

- Substantia nigra of C57BL/6J and CD-1 show no baseline differences in glial numbers
- Both mice show age and MPTP-induced gliosis in the substantia nigra pars compacta
- CD-1 nigra has lower levels of pro- and higher levels of anti-inflammatory cytokines
- Tilt of balance between pro- and anti-inflammatory cytokines begins at middle age
- Astrocytes and microglia show elongated mitochondria and intact ER upon MPTP-injection

## Introduction

Inflamm-aging refers to the alterations in the neuron-glia communication during aging, that result from a persistent functional decline in the immune system, characterized by a generalized increase in the pro-inflammatory markers (Franceschi et al., 2006). The system copes by releasing anti-inflammatory cytokines; a process termed as “anti inflamm-aging”. The imbalance between inflamm-aging and anti inflamm-aging associated processes, supported by the genetic make-up of the individual and environmental factors combine to trigger the onset or protect against age-related neurodegenerative diseases like Parkinson’s disease (PD). PD is characterized by a selective loss of dopaminergic (DA) neurons, primarily in the substantia nigra pars compacta (SNpc). The pattern of depletion in striatal dopamine and the dysfunction of the basal ganglia circuitry was considered as prognostic marker for cognitive dysfunction in drug naïve PD cases (Chung et al., 2018). It affects the natural gait while altering the cortico-striatal and cerebellar processing (Gilat et al., 2017). Non-motor symptoms like constipation, precede the motor symptoms by many years, thus suggesting a role for gut microbiota in the disease pathogenesis (Chaudhuri et al., 2006; Dutta et al 2019).

PD is characterized by T-cell infiltration (Galiano-Landeir et al., 2020; Subbarayan et al., 2020; Seo et al., 2020), microgliosis (Joers et al., 2017 review) and astrogliosis (Croisier and Graeber 2006), all of which affect the survival of substantia nigra neurons.Chronic neuroinflammation is considered to be a prototypical event preceding and accompanying neuronal dysfunction (Lee et al., 2009). It is marked by the presence of MHC class II activated microglia (Imamura et al 2003) and reactive astrocytes, direct participation of the adaptive immune system (Brochard et al., 2009; Harms et al., 2017); as well as increased synthesis of peripheral cytokines (Reale et al., 2009; Rydbirk et al., 2017); chemokines (Rastegar et al., 2019), inflammatory markers, reactive oxygen and nitrogen species(reviewed by Williams et al., 2019). Midbrain DA neurons express the receptors for cytokines such as tumor necrosis factor (TNF-α) and interleukin-1β (IL-1β),hinting at their sensitivity to these cytokines (Borrajoet al., 2014; Pang et al., 2015). Both neurons and glia express interferon-γ (IFN-γ) receptors and contribute to neuroinflammation (Hashioka et al 2010, Kim et al., 2013).

Epidemiological studies on PD propose prevalence and incidence rates of approximately 108-257/100,000 and 11-19/100,000 person year, respectively (Van Den Eeden et al., 2003) in Europe.North America reported 329-107/100,000 in Nebraska-USA (Strickland and Bertoni, 2004) and 224 per 100,000 person-years in persons above 65 years (Wirdefeldt et al., 2011). The incidence among Indians, Chinese and Malays were less compared to the Westerners (Gourie-Devi M 2014; Abbas et al., 2017). Thus, the white populations have significantly higher prevalence than non-whites; yet the underlying mechanisms are unclear. We demonstrated preservation of nigral neurons despite a non-logarithmic increase in α-synuclein with age, maintenance of GNDF receptors on substantia nigra neurons as well as other few neuroprotective factors in Asian Indians (Alladi et al., 2009; Alladi et al., 2010a; Alladi et al., 2010b). We also found age-related morphological transformation of astrocytes and microglia (Jyothi et al., 2015). Direct comparisons on human tissues between populations were not possible due to unavailability of tissues, which implored us to design an animal model to understand the mechanisms of differential prevalence.

Amongst the available small animal models for PD, a reliable recapitulation of PD is seen following administration of the neurotoxin 1-methyl-4-phenyl-1, 2, 3, 6-tetrahydropyridine (MPTP) in mice (Jackson-Lewis and Przedborski, 2007). In addition, different mice strains show varying responses to MPTP, for example, C57BL/6J mice is more sensitive while CD-1 white, BALB/c and Swiss Webster are resistant. Thus, the neurobiological attributes of these strains can be compared to understand the mechanisms of differential susceptibility and may be extrapolated to the human phenomenon of differential susceptibility to the disease. C57BL/6J dopaminergic neurons exposed to MPP^+^ (1-methyl-4-phenylpyridinium) demonstrated a 39% loss when cultured on C57BL/6J glia compared to 17% neuron loss when cultured on SWR/J glia (Smeyne et al., 2001). Thus, glia may modulate the strain-dependent susceptibility of mice to MPTP and comparison of responses of different mice strains may provide a platform to understand the population-based differences.

We have earlier reported that the CD-1 had more substantia nigra DA neurons than C57BL/6J and were better protected against MPTP (Vidyadhara et al., 2017). Soreq et al., (2017) reported that, astroglia show significant senescence-associated changes in gene expression patterns. Yet, the role of glia in aging, disease modulation, and differential vulnerability is not completely understood. In the present study we systematically investigated the responses of astroglia and microglia in the substantia nigra of the two different mice strains i.e. C57BL/6J and CD-1, in terms of aging and in association with inflammatory responses to MPTP. The outcome may be useful to understand the molecular mechanisms underlying the differences in prevalence of PD between the Caucasians and Asian-Indians.

## Materials and Methods

### Experimental Animals and MPTP-HCl Administration

We used C57BL/6J (MPTP-susceptible) and CD-1 (MPTP-resistant) mice at 15-17 week (young adults), 10-12 months (middle-aged) and 18-20 months (old/aged). All the experiments were conducted on male mice as there is a male preponderance in prevalence and incidence of PD (reviewed by Georgiev et al., 2017). They were housed under standard laboratory conditions of temperature 25° ± 2°C, 12 h light:12 h dark cycle with ad libitum access to food and water. The mice received four intraperitoneal injections of MPTP-hydrochloride (15 mg/kg/dose) in saline, at 2-h intervals. The controls received saline (Vidyadhara et al., 2017). The mice were sacrificed at days 1, 4 & 7 after MPTP-injection to assess the temporal profile of the cytokines. The other parameters were studied at 7 days post MPTP.

#### Tissue processing for immunohistochemistry (IHC)

Male mice anaesthetized using isoflurane were intracardially perfused with 0.9% heparinized saline followed by 4% ice-cold buffered paraformaldehyde (0.1M). The brains were removed and post-fixed for 48 hours at 4°C; cryoprotected in buffered sucrose grades (10%, 20% & 30%). 40μm thick serial coronal sections of midbrains were collected on gelatin coated slides.

#### Immunostaining

Two different series were used for Iba-1 and s100-β labeling (n=3-4/strain/age group/experimental condition); with a section periodicity of 1 in 6. Antigen was unmasked using sodium citrate buffer (pH-6) at 80°C for 20 min. Endogenous peroxidase was quenched using 0.1% hydrogen peroxide (H_2_O_2_) in 70% methanol (20 min in dark). The non-specific binding was blocked with 3% buffered bovine serum albumin (BSA) for 4 hr at room temperature (RT). The primary antibody exposure (0.1M PBS-TX; dilution 1:500; Table 1) lasted for 72 hours at 4°C in a closed chamber. Biotinylated secondary labeling (8 hr; dilution 1:100, Vector labs, USA; Table 1) was followed by tertiary labeling with avidin-biotin complex (Vector labs, USA; 4 hr at RT). The staining was visualized with 0.05% DAB (3’-3’-diaminobenzidine) and 0.01% nickel sulphate in 0.1M acetate imidazole buffer (pH 7.4) containing 0.01% H_2_O_2_ resulting in a black colored reaction. For the second antibody, similar procedure was followed (Table 1) except that the chromation was performed exclusively with DAB, resulting in brown colored profiles [(TH (brown) and Iba-1/s100β black)]. Negative controls were processed without adding primary antibody. The sections were mounted with DPX following alcohol-based dehydration.

**TABLE 1:** Details of Primary and Secondary antibodies used for immunochemistry.

| Staining modality, primary antibody and source |  | Dilution and incubation time | Secondary antibody and ABC kits | Dilution and incubation time |
| --- | --- | --- | --- | --- |
| Immuno-<br>peroxidase<br>Staining | Anti-Goat Iba-1,<br>Abcam, UK | 1:800<br>72 hr at 4°C | Goat Elite ABC kit,<br>Vector laboratories,<br>USA | 1:200, 4 hr at<br>RT |
| | Anti-Mouse S100β,<br>Sigma –Aldrich | 1:800<br>72 hr at 4°C | MOM* ABC kit,<br>Vector<br>laboratories,USA | 1:500, 3 hr at<br>RT |
|  | Anti-Rabbit TH,<br>Santa Cruz, USA | 1:1000<br>72 hr at 4°C | Rabbit Elite ABC<br>kit,<br>Vector<br>laboratories,USA | 1:200, 4 hr at<br>RT |
| Immuno-<br>fluorescence | Anti-Goat Iba-1,<br>Abcam UK | 1:800<br>72 hr at 4°C | FITC, Sigma<br>Aldrich,USA | 1; 200 6-8 hr at<br>4°C |
|  | Anti-Rabbit<br>Fractalkine, Abcam<br>UK | 1:500<br>72 hr at 4°C | Cy3, Sigma Aldrich,<br>USA | 1; 200 6-8 hr at<br>4°C |

MOM\*: Mouse-on-mouse kits

For immunofluorescence labeling, following the incubation with primary antibody, the sections were incubated in an appropriate fluorescently labeled secondary antibodies (Sigma-Aldrich, USA). The sections were then mounted using Vectashield hard set mounting medium (Vector Laboratories, USA) and images were captured using a laser scanning confocal microscope (DMIRE-TCS Leica Microsystems, Germany).

#### Stereological quantification of cells

The s100β and iba-1immunoreactive (ir) cells in SNpc were quantified using optical fractionator method with an Olympus BX61 Microscope (Olympus Microscopes, Japan) equipped with Stereo Investigator software version 8.1 (Micro-brightfield Inc., Colchester, USA). The SNpc in TH-ir midbrain sections was delineated on both the sides under 10X (Paxinos, 2013). Pilot studies determined the grid interval and counting frame size. Cells were counted in every sixth section (6-8 sections/animal) under 100X, with a grid interval of 10000μm^2^ (x=100μm, y=100μm) and counting frame of size 6400μm^2^ (x=80μm, y=80μm) as per our earlier report (Vidyadhara et al., 2017). The absolute numbers were derived and coefficient of error was determined according to Gundersen et al., (1999).

#### Immunoblotting

The mice (n=5/strain/age group/experimental condition) were sacrificed by cervical dislocation and the midbrains were snap frozen in liquid nitrogen and stored at −80°C till further use. The brains were thawed to −20°C on a cryostat and 5μm thick sections of SN were solubilized in 100 ul of mammalian lysis buffer 10% protease inhibitor cocktail (Sigma–Aldrich, USA). Following sonication (Q sonica, India) and centrifugation at 12000g (4°C) the protein concentration in the supernatant was assayed by Bradford’s method and 60ug of protein/sample was electrophoresed (Bio-Rad, USA) on a 5%/10% (loading/separating) denaturing gel. The proteins were transferred onto a buffered poly-vinilidine di-fluoride (PVDF) membrane (Millipore, Germany). The non-specific staining was blocked by 5% skimmed milk protein (1X TBST;4hr) followed by overnight incubation with primary antibody solution (Table 2.). This was followed by incubation with appropriate HRP-conjugated secondary antibody for 2hr (Table 2.). The band was detected using chemi-luminescent substrate for HRP (Super Signal West Pico, USA) using a geldoc apparatus (Syngene International Ltd., India) and quantified using Image J 1.48 v program (Vidyadhara et al., 2017). B-actin was used as a loading control and each protein concentration was normalized against it.

**TABLE 2:** Details of primary and secondary antibodies used for immunoblotting.

| **Primary antibody** | **Dilution and incubation time** | **Secondary antibody** | **Dilution and incubation time** |
| --- | --- | --- | --- |
| Anti iNOS, Rabbit, Abcam USA | 1:1500,<br>8 hr at 4°C | Anti Rabbit HRP-Conjugated secondary antibody, Merck | 1:2000,<br>2 hr at RT |
| Anti Fractalkine, Rabbit, Abcam USA | 1:800,<br>8 hr at 4°C | Anti Rabbit HRP-Conjugated secondary antibody, Merck | 1:2000,<br>2 hr at RT |
| Anti β-actin, Mouse, Sigma Aldrich | 1:2000,<br>8 hr at 4°C | Anti Mouse HRP-Conjugated secondary antibody, Merck | 1:2000,<br>2 hr at RT |

#### Enzyme-linked immunosorbent assay (ELISA)

Protein lysates were obtained from ventral midbrains (n=4-5/strain/age-group/time point/experimental condition) of post-MPTP day (d)1, d4 and d7 mice. Both pro-inflammatory (TNF-α, IL-6, IL-1β) and anti-inflammatory cytokines (TGF-β, IL-4, 1L-10) were evaluated as per the manufacturer’s instructions (Ray Biotech, Inc, USA). MAO-A and MAO-B kits were obtained from Cloud-clone Corp, (Wuhan, China) and HO-1 ELISA kit was obtained from Abcam UK (mouse Simple Step, ab204524). Briefly 100μl of standards/samples were added to each well except blanks/standards (2.5 hr, 37°C). Thereafter, 100μl of biotinylated secondary antibody was added (1hr, RT) with gentle shaking followed by streptavidin (100μl). The chromation was achieved with TMB (3,3’,5,5’-tetramethylbenzidine) one-step substrate reagent. The reaction was stopped with 50μl stop solution/well and the absorbance was measured immediately at 450nm (TECAN, Austria). All standards and samples were run in duplicates.

### Electron microscopy

The mice (n= 3/strain/age group/experimental condition) were anesthetized with isoflurane and transcardially perfused with a buffered mixture of 2.5% glutaraldehyde and 2% PFA (in 0.1M phosphate buffer, pH 7.4). The dissected substantia nigra was cut into smaller pieces, post-fixed with 1% osmium tetroxide at RT and dehydrated in grades of ethyl alcohol followed by clearing in propylene oxide. Infiltration (1:1 mixture of araldite and propylene oxide, overnight at RT, on a rotator) was followed by exposure to pure araldite for 4.5h. The resin embedded tissues (flat embedding molds) were allowed to polymerize (60°C, 48 hr). 1μm semi thin sections were stained with 1% toluidine, blue to verify the region of interest and 60nm ultrathin sections (Leica, Ultramicrotome, Austria) collected on copper grids were stained using saturated uranyl acetate for 1 hr and 0.2% lead citrate (5-7 min). The washed and air-dried sections were examined under Transmission Electron Microscope (FEI, TECNAI G2 Spirit BioTWIN, Netherlands) and images were captured using Mega View-III CCD camera (Shruthi et al., 2017).

#### Statistics

The stereology-based data for quantification of astrocytes and microglia was analysed by 1 Way ANOVA followed by Bonferroni post hoc analysis. The ELISA related data was analyzed using Kruskal Wallis test and Mann-Whitney U tests and represented as line graphs with median values with upper range for each strain and time point. The other observations were analyzed using two-way ANOVA followed by Tukey’s post hoc test using SSPS software and GraphPad Prism software (GraphPad Software version 6.01Inc, USA). A p–value <0.05 was considered as statistically significant. The data was expressed as mean ± standard error of mean (mean ± SEM).

## RESULTS

### I. Differences in number of glia

#### A. Microglia

The strain-specific differences in the neuronal numbers of substantia nigra noted earlier (Vidyadhara et al., 2017) prompted us to verify if the differences extended to baseline numbers of glia. Iba-1 stained the microglia well (fig. 1A&B; arrows), including their processes and were easily discriminated from neurons (arrowheads) in the vicinity. Based on unbiased stereology, CD-1 nigra had relatively fewer Iba-1 ir cells across aging although not statistically significant (fig.1C; 18.2% at young age; 14% at middle-age and 8.5% at old age). In response to MPTP, an increase in microglia was noted in old animals of both strains [C57BL/6J MPTP (young vs. middle-aged ## p<0.01, young vs. old # p<0.05), CD-1 MPTP (young vs. old $ p<0.05)].

**Figure1:**
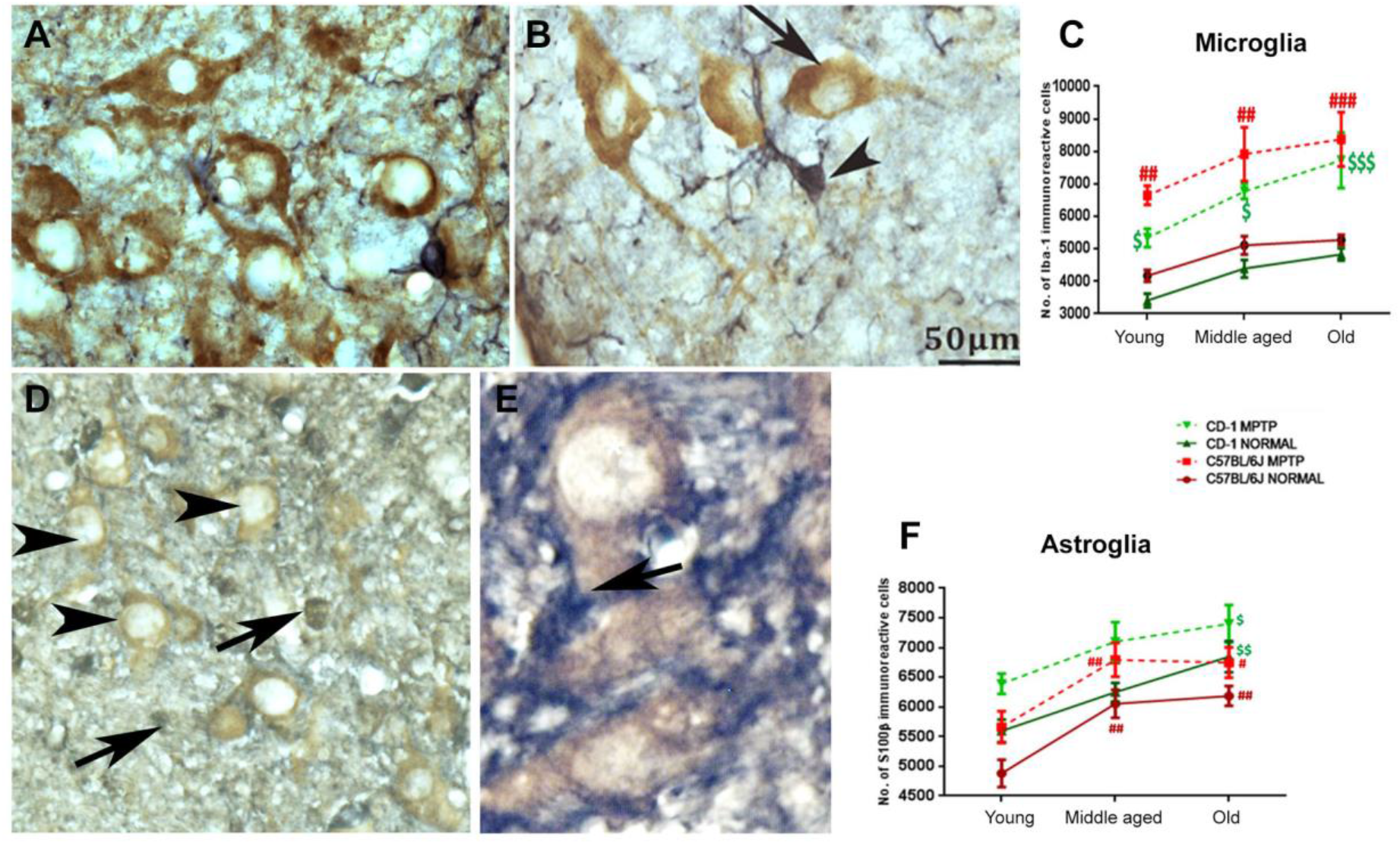
Differences in numbers of microglia and astrocytesin the two strains following MPTP administration. **A&B)** Representative DAB-stained photomicrographs of Iba-1 ir microglia (black) and TH positive DA neurons (brown) within the SNpc of 18-month-old C57BL/6J (A) and CD-1 mice (B).The line graph(C) shows no significant interstrain differences in control mice. Note the gradual increase in the microglial numbers during aging in both strains. Note the significant increase in the number of microglia at all ages after MPTP injection [Young (C57BL/6J normal vs. C57BL/6J MPTP ## p<0.01, CD-1 normal vs. CD-1 MPTP # p<0.05); middle-aged (C57BL/6J normal vs. C57BL/6J MPTP ## p<0.01, CD-1 normal vs. CD-1 MPTP $$ p<0.01); old (C57BL/6J normal vs. C57BL/6J MPTP ###p<0.001, CD-1 normal vs. CD-1 MPTP $$$ p<0.001),(* between strains, #within C57BL/6J,^$^within CD-1). Scale bar: 50 μm. **Astrocytes and susceptibility:** Representative DAB-stained photomicrographs of s100β-ir astrocytes (black) and TH immunopositive DA neurons (brown) in SNpc of 18 months old C57Bl/6J (**D)** and CD-1 **(E)** mice. The line graph (**F)** a significant increase in the numbers of astrocytes during aging in both strains and in response to MPTP. C57BL/6J normal (young vs. middle-aged ## p<0.01, young vs. old ## p<0.01), C57BL/6J MPTP (young vs. middle-aged ## p<0.01, young vs. old # p<0.05), CD-1 normal (young vs. old $$ p<0.01) CD-1 MPTP (young vs. old $ p<0.05) (difference * between strains, #within C57BL/6J,^$^within CD-1). Scale bar: 50 μm.

#### B. Astrocytes

The astrocytes appeared much darkly stained (fig. 1D&E; arrows) compared to the TH ir dopaminergic neurons (arrowheads). The control CD-1 mice substantia nigra had moderately more s100β positive astrocytes (fig. 1B) than the C57BL/6J at young (fig.1C&D, 14.68%) and old age (fig. 1C&D, 10.67%). Under normal conditions, in C57BL/6J, the numbers increased significantly from youth to middle-age (C57BL/6J young vs. middle-aged ^##^p<0.01; 24.06%) to stabilize later (C57BL/6J young vs. old ## p<0.01; 26.75%). In contrast, the CD-1 substantia nigra showed a gradual age-related increase in the astrocytes (CD-1, young v/s middle age, 11.64%; young vs. old $$ p<0.01; 22.32%). MPTP caused a significant increase in the numbers at all ages, in both the strains; notably higher in the young mice (C57BL/6J control v/s MPTP 20%; CD-1 control vs. MPTP, 15.8%).

### II. Cytokine expression in the ventral midbrain

Increase in the numbers of microglia and astrocytes in response to MPTP led us to hypothesize that gliosis may induce neuroinflammation. We therefore estimated the levels of pro and anti-inflammatory cytokines.

#### A. Pro-inflammatory cytokine TNF-α was relatively higher in C57BL/6J

The baseline TNF-α expression peaked at middle-age in the C57BL/6J (fig 2. top panel; normal aging; ^####^ p<0.0001; young vs. middle age); which was also significantly more than CD-1 (*p=0.032; C57BL/6J vs. CD-1). At old age too, the differences persisted, with higher levels in C57BL/6J (*p=0.016; C57BL/6J vs. CD-1). MPTP caused an up-regulation in the middle-aged C57BL/6J at d1(C57BL/6J control vs. post-MPTP d1, ^#^p=0.032) and remained higher than CD-1 (C57BL/6J vs. CD-1, **p=0.008). In CD-1 the differences were noted at d4 (control vs. d4 post-MPTP; ^$$$^p=0.001). The levels were overall higher in C57BL/6J, both at d1 and d7 (C57BL/6J vs CD-1; d1,**p=0.008 and d7,*p=0.016). At old age, C57BL/6J showed a significant increase at d1 (control vs. post-MPTP d1, ^#^p=0.032). Further at d7, the differences were evident between the strains (C57BL/6J vs.CD-1 *p = 0.032).

**Figure 2:**
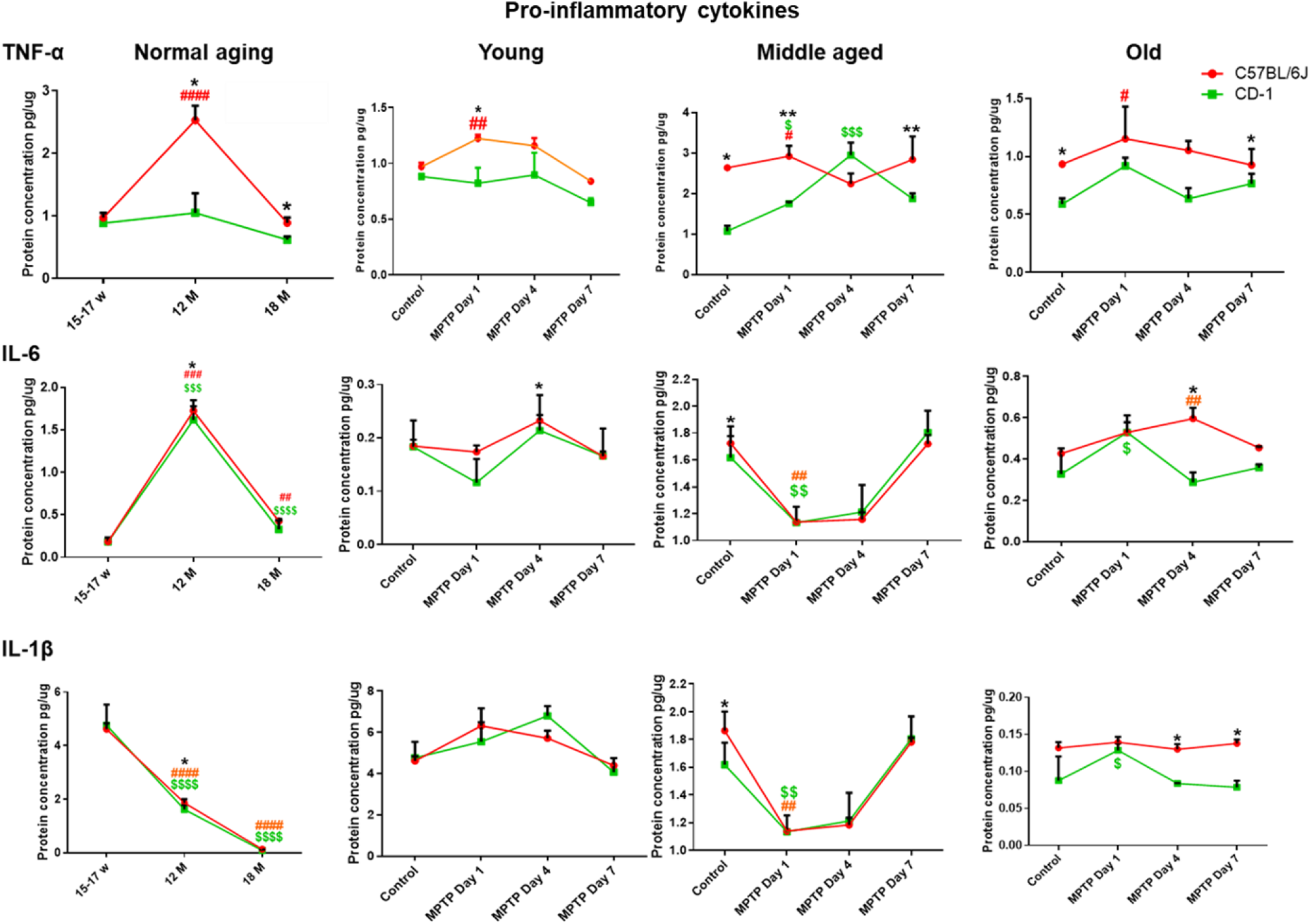
Levels of Pro-inflammatory cytokines: Line graphs representing different strains at different age periods and in response to MPTP. The top horizontal panel pertains to TNF-α, middle to IL-6 and bottom panel to IL-1β. Note, differences are depicted as * = between strains, ^#^ = within C57BL/6J, ^$^= within CD-1. **A) Higher pro-inflammatory cytokine levels (TNF-α) in C57BL/6J:** Note higher levels in C57BL/6J than CD-1 at middle (*p=0.032) and old age (*p=0.016). C57BL/6J shows peak TNF-α expression at middle-age (^####^ p<0.0001; young vs. middle age). At middle-age MPTP induced increase in TNF-α in C57BL/6J (control vs. post-MPTP d1, ^#^p=0.032) and CD-1 (control vs. d4 post-MPTP; ^$$$^p=0.001). The levels were higher in C57BL/6J at d1 and d7 (C57BL/6J vs CD-1; d1,**p=0.008 and d7,*p=0.016). Old C57BL/6J showed more expression at d1 (control vs. post-MPTP d1, #p=0.032). Note the differences between strains at d7 (C57BL/6J vs.CD-1 *p = 0.032). **B) Middle-aged animals showed significant increase in IL-6:** Note the higher IL-6 expression at middle age (C57BL/6J young vs. middle-age ^###^p=0.0007; CD-1 young vs. middle-aged ^$$$^p<0.0001). C57BL/6J showed higher expression than CD-1 (*p = 0.032). Note the reduction at old age (C57BL/6J middle-age vs. old ^##^p=0.0010; CD-1 middle-age vs. old ^$$$$^p<0.0001). At 1d, a reduction is seen at middle age, in response to MPTP in C57BL/6J (^##^p=0.008) and CD-1 (^$$^p=0.008). MPTP caused an upregulation in old CD-1 at d1 (control vs. d1, ^$^p=0.016); while at d4 in old C57BL/6J (control vs. d4 ^##^p=0.008), which is higher than CD-1 (C57BL/6J vs. CD-1 post-MPTP d4*p=0.032). **C) IL-1β is down regulated during normal aging:** Note the age associated reduction in IL-1β in both strains (C57BL/6J young vs. middle-aged ^####^p<0.0001; CD-1 young vs. middle-aged ^$$$$^p<0.0001; C57BL/6J middle-aged vs. old ^####^ p<0.001; CD-1 middle-aged vs. old ^$$$$^p<0.001). Inter-strain differences are present at middle age, with higher levels in C57BL/6J (C57BL/6J vs. CD-1; *p=0.032). Note the reduction upon MPTP in middle aged C57BL/6J (control vs. post-MPTP d1^##^p=0.01) as well as CD-1 (control vs. post-MPTP d1; ^$$^p=0.01). Note the increase in old CD-1 at d1 (control vs. post-MPTP d1; CD-1 ^$^p=0.032). Post-MPTP at d4 and d7 the levels remain relatively higher in C57BL/6J (C57BL/6J vs. CD-1, *p=0.032).

##### IL-6 and IL-1β show largely similar patterns and MPTP responses in both strains

Both the strains showed a significantly higher IL-6 expression at middle-age (fig 2; C57BL/6J young vs. middle-age ^###^p=0.0007; CD-1 young vs. middle-aged ^$$$^p<0.0001). Between the strains C57BL/6J showed higher expression (C57BL/6J vs. CD-1, *p = 0.032). Thereafter a significant reduction ensued at old age in both the strains (fig 2; C57BL/6J middle-age vs. old ##p=0.0010; CD-1 middle-age vs. old ^$$$$^p<0.0001). At middle age, in response to MPTP, both C57BL/6J and CD-1 showed a notable reduction at d1 (^##^p=0.008; ^$$^p=0.008) and were almost restored to the baseline at d7. At old-age the MPTP-induced upregulation was appreciated at d1, in CD-1 (^$^p=0.016, control vs. d1; CD-1). The old C57BL/6J showed an increase at d4 (post-MPTP d4 ^##^p=0.008), and was relatively more than CD-1 at that time point (d4,*p=0.032).

IL-1β showed a significant reduction with aging in both strains (C57BL/6J young vs. middle-aged ^####^p<0.0001; CD-1 young vs. middle-aged ^$$$$^p<0.0001; C57BL/6J middle-aged vs. old ^####^ p<0.001; CD-1 middle-aged vs. old ^$$$$^ p<0.001). Baseline differences between strains were noted only at middle age, with higher expression in C57BL/6J (C57BL/6J vs. CD-1; *p=0.032). The MPTP-injected middle aged C57BL/6J (control vs. post-MPTP d1^##^p=0.01) as well as CD-1 (control vs. post-MPTP d1; ^$$^p=0.01) showed an acute decrease at d1.

The old CD-1 mice strains showed an augmented response at d1 (control vs. post-MPTP d1; CD-1 ^$^p=0.032). Post-MPTP at d4 and d7 the levels remain relatively higher in C57BL/6J vis-à-vis the CD-1 (*p=0.032; C57BL/6J vs. CD-1).

##### Levels of anti-inflammatory cytokine TGF-β are relatively lower in C57BL/6J

The basal level of TGF-β was significantly lower in C57BL/6J at all the age periods studied [(fig 3; top panel; C57BL/6J vs. CD-1; young **p=0.008), middle-aged (**p=0.008), old (**p=0.008)]. In response to MPTP, middle aged C57BL/6J showed an increase in expression (C57BL/6J control vs. post-MPTP d1, #p<0.032). Middle-aged CD-1 showed more expression than C57BL/6J in normal conditions (C57BL/6J vs. CD-1 **p=0.08). Moreover, in response to MPTP, the levels remained higher at all the days studied (C57BL6/J v/s CD-1 d1 *p=0.036; d4 *p=0.029; d7 *p=0.016). At old age too, the CD-1 showed more expression than C57BL/6J, in normal conditions (C57BL/6J vs. CD-1 **p=0.008). In response to MPTP, at d1, despite the decrease in CD-1 (control vs. d1 ^$^p=0.016), the levels were comparatively higher than the C57BL/6J (C57BL/6J vs. CD-1 *p=0.016); as also at d7 (C57BL/6J vs. CD-1 *p=0.05).

**Figure 3:**
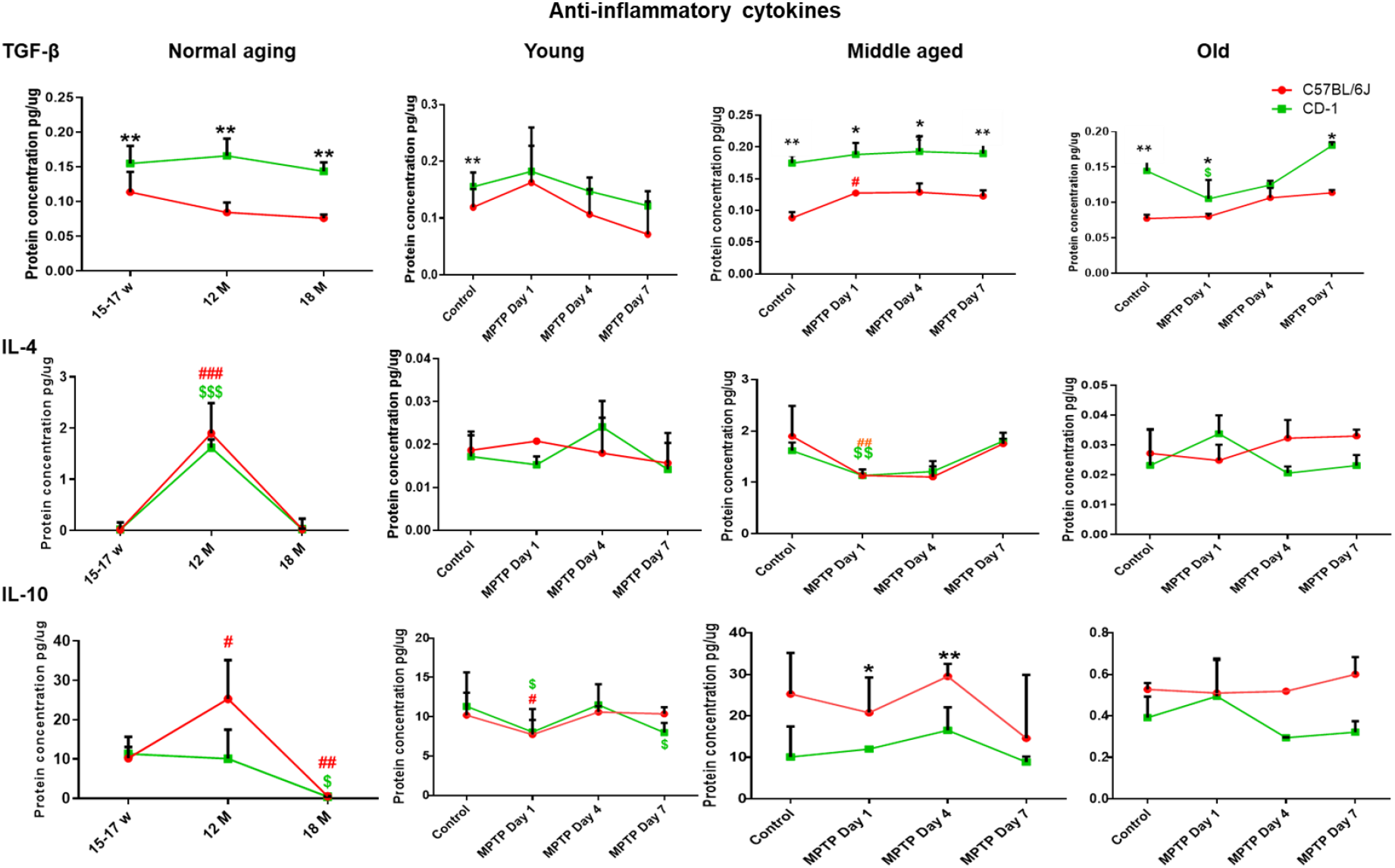
Levels of anti-inflammatory cytokines: Line graphs representing different strains at different age periods and in response to MPTP. The top horizontal panel pertains to TGF-β, middle to IL-4 and bottom panel to IL-10. Note, the differences are depicted as * = between strains, ^#^ = within C57BL/6J, ^$^ =within CD-1. **A) Lower levels of anti-inflammatory cytokine TGF-β in C57BL/6J:** Note the lower basal TGF-β levels in C57BL/6J through aging (C57BL/6J vs. CD-1; young **p=0.008; middle-aged **p=0.008; old **p=0.008). MPTP caused upregulation in middle aged C57BL/6J at d1 (C57BL/6J control vs. d1 ^#^p<0.032). However, overall expression was higher in CD-1 (C57BL6/J v/s CD-1 d1 *p=0.036; d4 *p=0.029; d7 *p=0.016). Note the decrease in old CD-1 at d1, upon MPTP, (control vs. d1 ^$^p=0.016) and yet higher levels than the C57BL/6J (d1 C57BL/6J vs. CD-1 *p=0.016; d7 C57BL/6J vs. CD-1 *p=0.05). **B) Increase in the basal level expression of IL-4 in middle-aged animals:** Both the strains show an increase in IL-4 at middle-age (C57BL/6J young vs. middle age ^###^p=0.0010; CD-1 young vs. middle age ^$$$^p=0.0009). Note the decrease at old age (C57BL/6J middle age vs. old ^###^p=0.0010; CD-1 middle age vs. old ^$$$^p=0.0002). Note the MPTP-induced reduction at d1 in both strains (C57BL/6J##p=0.008; CD-1 ^$$^p=0.008), at middle age. Old age observations are comparable between strains. **C) Compensatory response by IL-10 in C57BL/6J**: Note the increase in levels in C57BL/6J from young to middle age (C57BL/6J young vs. middle age ^#^p=0.0125) and a decrease at old age (C57BL/6J middle age vs. old ^#^p=0.0033). The decrease in CD-1 is seen between middle and old age (CD-1 ^$^p=0.0317). Upon MPTP, young C57BL/6J (^#^p=0.032) and CD-1 (^$^;p=0.032) show a reduction at d1. Note, the slight increase at d4 and a reduction at d7 in CD-1 (^$^p<0.033). At middle age, MPTP-induced inter-strain differences are appreciable at d1 and d4, with higher expression in C57BL/6J (C57BL/6J vs. CD-1; d1 *p=0.016; d4 **p=0.008). Note the non-significant differences at old age.

Both the strains showed a significant increase in IL-4 expression at middle-age (Fig 3; C57BL/6J young vs. middle age ^###^p=0.0010; CD-1 young vs. middle age ^$$$^p=0.0009). Thereafter a decrease was noted at old age (C57BL/6J middle age vs. old ^###^p=0.0010; CD-1 middle age vs. old ^$$$^p=0.0002). At middle age, MPTP elicited a reduction at d1 in both strains (C57BL/6J^##^p=0.008; CD-1 ^$$^p=0.008).

#### B. Anti-inflammatory cytokine IL10 levels are relatively higher in middle aged C57BL/6J

The control C57BL/6J showed an increase in expression from young to middle age (C57BL/6J young vs. middle age ^#^p=0.0125) and a much appreciable decrease by old age (C57BL/6J middle age vs. old ^#^p=0.0033); whereas the CD-1 showed a decrease from middle to old age (CD-1 middle age vs. old, ^$^p=0.0317).

MPTP elicited strain specific responses. At young age, both C57BL/6J (^#^p=0.032) and CD-1 (^$^p=0.032) showed a reduction in IL-10 expression in response to MPTP at d1. The expression increased slightly at d4 only to reduce further in CD-1 at d7 (^$^p<0.033). At middle-age both strains showed differences in baseline expression, which were however not statistically significant. The MPTP-induced inter-strain differences were noted, with higher expression in C57BL/6J at d1 and persisted till d4 (C57BL/6J vs. CD-1; d1 *p=0.016; d4 **p=0.008). Further, the old C57BL/6J showed mildly higher expression in response to MPTP at d4 and d7, however the differences did not reach significance.

### III: Differences in enzyme expression

#### A: MAO-A and MAO-B levels

Monoamine oxidase (MAO) is a vital enzyme that metabolizes biogenic amines like noradrenaline, adrenaline, dopamine etc. MAO-B present in glia is responsible for the breakdown of MPTP to MPP^+^. An interesting comparison between controls and PD patients showed that the PD striatum harbors significantly higher levels of MAO-B compared to controls; whereas the substantia nigra of both controls and PD showed no differences (Tong et al., 2017). We therefore conducted the MAO-B enzyme analyses in the striatal specimens. MAO-A positive punctae were localized to the cytoplasm (Fig.4A1, red) of the TH immunoreactive DA neurons (Fig.4A2) as also in the neuropil (Fig.4A3 pink coloration in neurons and red in the neuropil). The basal levels as measured by immunoblotting were higher in the substantia nigra of young CD-1 (C57BL/6J vs. CD-1 *p<0.05), however middle-age saw a peak in C57BL/6J (young vs. middle aged ^###^p <0.001). In both strains, the levels reduced at old age (fig 4B, C57BL/6J middle aged vs old *###*p<0.001; CD-1 middle aged vs. old ^$$^ p>0.01). MPTP induced a notable increase in the young C57BL/6J; although both strains showed moderate enhancement at all ages (C57BL/6J control vs. MPTP ^###^p<0.001).

**Figure 4:**
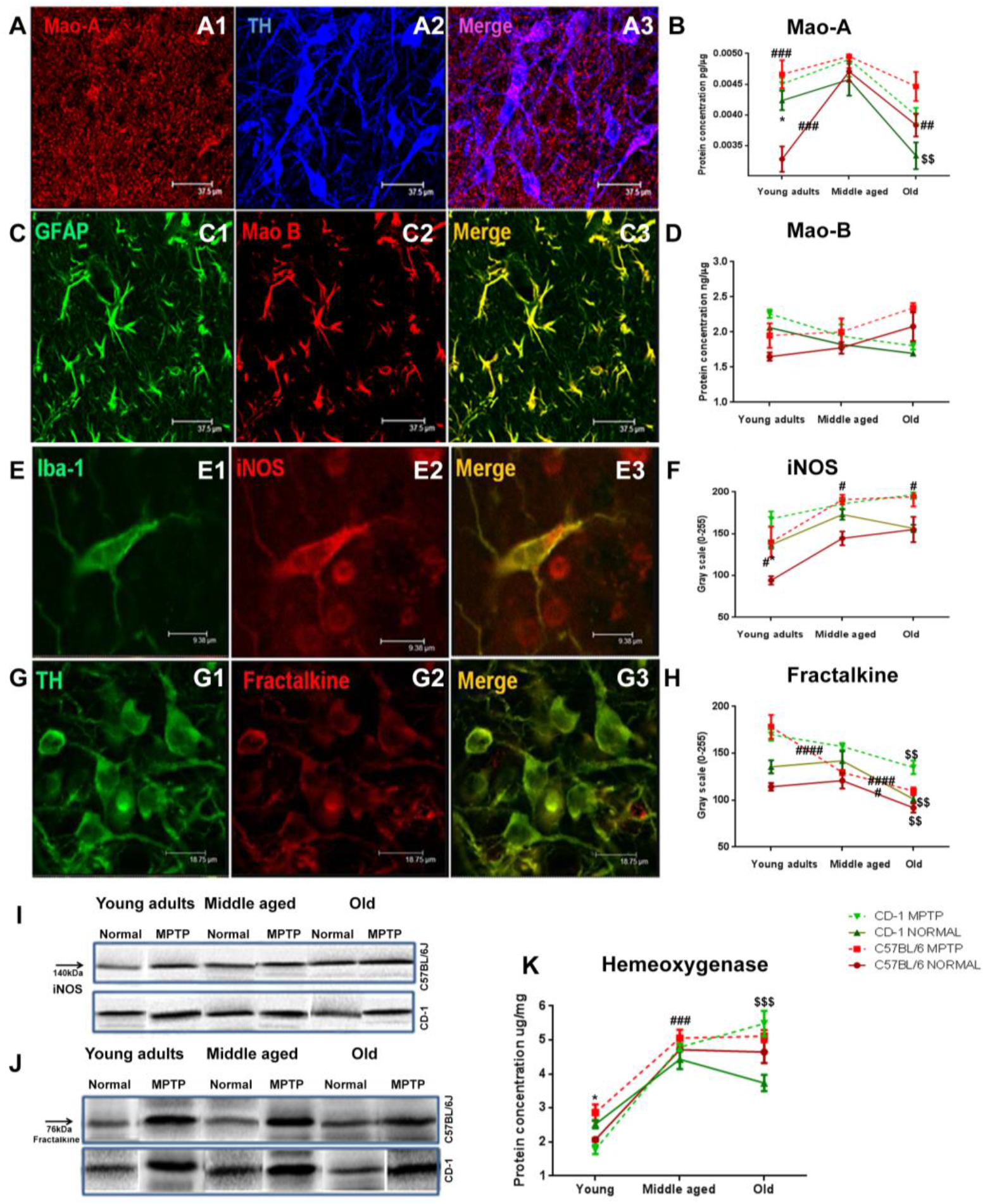
Essential Enzymes are differentially modulated in a strain specific manner: **A)** Representative photomicrographs of DA neurons (A1, TH, blue; cytoplasmic) co-labeled with MAO-A (A2, red; punctate staining), and merge showing presence of MAO-A in dopaminergic neurons in SNpc (A3; pink). Note the punctate staining of MAO-A vis-a-vis the pan cytoplasmic staining of TH. **B)** Representative immunofluoresence photomicrographs of astrocytes (GFAP, green, B1) co-labeled with MAO-B (red, B2) in striatum and the merge showing glial localization of MAO-B (B3, yellowish green). **B&D).** The basal levels of MAO-A and MAO-B are higher in CD-1 (MAO-A C57BL/6J vs. CD-1 * p<0.05). Note the significant upregulation of MAO-A level in middle-aged C57BL/6J (young vs. middle-aged ### p<0.001). Aging increased the MAO-B levels in C57BL/6J whereas CD-1 showed a down regulation. MPTP enhanced MAO-A and MAO-B levels in both strains. (Difference * between strains, ^#^within C57BL/6J, ^$^within CD-1).**E)** Representative immunofluorescence photomicrographs of microglia (E1, Iba-1, green) co-labeled with iNOS (E2, red) and their co-labeling (E3, merge, yellowish orange) in SNpc. Note the presence of some INOS ir cells that were not Iba-1 ir. (E3, red). **F).** Histograms show that basal iNOS level was significantly higher in CD-1 mice in young adults (C57BL/6J vs. CD-1 * p<0.05; I) **I)** Western blot showing a single band of iNOS at 140kDa. Note the gradual increase at middle-age in both strains. MPTP caused a strain independent augmentation in all age groups (middle-aged C57BL/6J control vs. MPTP# p<0.05, old C57BL/6J control vs. MPTP# p<0.05, old CD-1 control vs. MPTP$ p<0.05. **G)** Representative immunofluorescence photomicrographs of DA neurons (G1, TH, green) co-labelled with fractalkine (G2, red) and their co-labeling (G3, merge, yellow-green) in SNpc. **H)** histograms show higher levels in CD-1 at all ages. Both strains show a gradual decrease with age (C57BL/6J middle-age vs. old ^#^p<0.05, CD-1 middle-age vs. old ^$$^p<0.01). In C57BL/6J, the response to MPTP at middle and old age was lesser (young vs. middle-aged^####^p<0.0001, middle-age vs. old ^####^p<0.0001).CD-1 show a slow gradual reduction. (Difference **\*** between strains, ^#^within C57BL/6J, ^$^within CD-1). **J).** Representative Western blot bands of fractalkine showing single band at 76KDa in different study groups. **K)** Compensatory response by Hemeoxygenase-1:Histograms showing lower levels in young C57BL/6J, whereas CD-1 shows stabilization at old age (C57BL/6J young adult vs. middle-aged ^####^ p<0.0001, CD-1 mice young adult vs. middle-aged $$$ p<0.001). Both strains show increased expression after MPTP (C57BL/6J vs CD-1, young *p<0.05; CD-1 old control vs. MPTP$$$ p<0.001), (Difference *between strains, #within C57BL/6J, $ within CD-1).

The GFAP expressing glia (fig.4B1, green) showed MAO-B immunoreactivity (fig.4B2, red fig.4B3 merge, yellowish green). MAO-B levels were also higher in young CD-1 than C57BL/6J (fig. 4fD). An age-associated gradual down-regulation in CD-1 was contrasted by a gradual increase in C57BL/6J, although statistically significant. In summary, MPTP administration up regulated both MAO-A and MAO-B level in both strains

#### B: Inducible nitric oxide synthase (iNOS)

Chhor et al., (2013) reported that reactive microglia in the degenerating SN led to the secretion of iNOS. Accordingly, in our specimens too Iba-1 immunopositive microglia (fig. 4E1; green) were primarily noted to express iNOS (fig. 4E2; red) in addition to some non-microglial cells (fig. 5E3 merge; red). The antibody showed a single band of 140 K Da (fig. 4I). Young CD-1 had significantly higher basal iNOS (fig. 4F; C57BL/6J vs. CD-1 *p<0.05). At middle-age both strains showed a moderate increase in expression. With aging, the CD-1 mice showed mild decrease in iNOS expression whereas C57BL/6J maintained the levels attained at middle age. MPTP caused iNOS augmentation across ages in both strains (middle-aged C57BL/6J control vs. MPTP #p<0.05, old C57BL/6J control vs. MPTP #p<0.05, old CD-1 control vs. MPTP $p<0.05).

**Figure 5:**
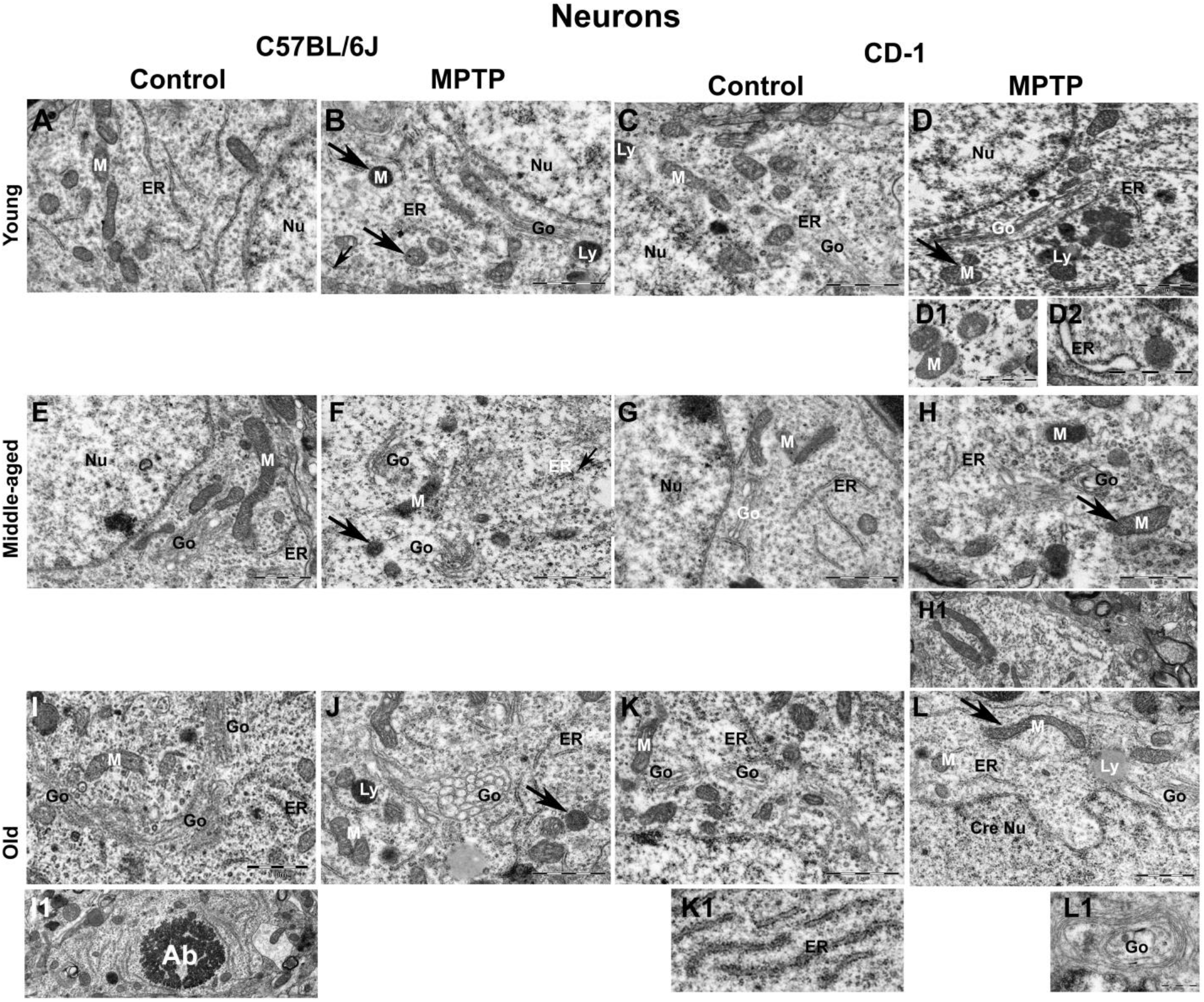
Age-related and MPTP-induced ultrastructural changes in substantia nigra neurons. Representative electron micrographs of substantia nigra neurons at different ages and in response to MPTP. Note that the mitochondria of the elderly C57BL/6J were relatively shrunken (compare A,E&I). MPTP caused mitochondrial shrinkage at all ages (compare A&B and E&F, I&J; ‘M’ arrows). In CD-1 the mitochondrial size was fairly preserved with age (compare C&D, D1 and G&H, K&L; ‘M’ arrows) or longer in response to MPTP (H, H1 and L, arrow). Note the shortening of ER strands with age (compare A,E&I; ER) and in response to MPTP in C57BL/6J (compare A&B and E&F, I&J; ER). Note the dilated ER in CD-1 neurons upon MPTP in young (D2) and at middle-age (ER, H & H1). Note the ER arrays in aged CD-1 (K1) and presence of many Golgi units in middle aged and old mice of both strains (‘Go’ in I&K). MPTP-administered aged C57BL/6J have circular Golgi with bloated saccules (J&L1; ‘Go’). Older C57BL/6J have apoptotic bodies (I1; ‘Ab’) and CD-1 has crenellated neuronal nucleus. Note several lysosomes in young MPTP injected CD-1 against few in C57BL/6J (B&D ‘Ly’). The scale bar for A=B, and others are specified for each photomicrograph.

#### C: Fractalkine

Fractalkine (CX3CL1) is an enzyme synthesized by neurons, and it suppresses microglial activation (Thome et al., 2015). In our observation, it was localized to the DA neuronal cytoplasm (fig.4G1-G3). The antibody showed a band at 140 KDa (fig. 4J). The CD-1 substantia nigra had moderately high levels at all the ages, compared to C57BL/6J. A decrease was evident in both strains after middle age (Fig. 4H, C57BL/6J middle-age vs. old # p<0.05, CD-1 middle-age vs. old $$ p<0.01). MPTP elicited an increase in both the strains, but more appreciably in the young (C57BL/6J, young adults vs. middle-aged #### p<0.0001, middle-aged vs. old #### p<0.0001).

#### D: HO-1

This family of enzymes, along with NADPH cytochrome P450 reductase, modulates the catabolism of cellular heme to biliverdin, carbon monoxide (CO) and free ferrous ironin brain (Ryter and Tyrrell, 2000).The young CD-1 midbrains showed mildly higher levels of HO-1which reduced following MPTP. Both strains showed a significant up-regulation at middle-age that persisted till old age (fig. 4k, C57BL/6J young vs. middle-aged ^####^p<0.001, CD-1 young vs. middle-aged $$$p<0.001). Aged mice of both strains showed an increase in HO-1 expression in response to MPTP; moderate in C57BL/6J; significant in CD-1 (CD-1 control vs. MPTP $$$p<0.001), suggesting a compensatory induction.

### IV: Age-related and MPTP-induced ultrastructural changes

#### A: Effects on Substantia Nigra Neurons

The mitochondria of the young and middle aged C57BL/6J were relatively larger than those of the elderly (fig. 5A,E&I). MPTP induced mitochondrial shrinkage at all ages (fig.5 compare A&B and E&F, I&J; ‘M’ arrows). Shrunken albeit intact mitochondria were evident, in the MPTP injected young CD-1 mice (fig. 5 D). However, at middle and old age they were preserved (fig. 5 compare C&D and G&H, K&L; ‘M’ arrows) or even elongated in response to MPTP (fig. 5 H, H1 and L, arrow). The endoplasmic reticular (ER) strands shortened with age and in response to MPTP in C57BL/6J (fig. 5, compare A&B and E&F, I&J; ER). Whilst MPTP induced ER dilatation in the young (fig. 5; D2) and middle-aged CD-1 mice neurons (fig. 5H; ER). The normal aged CD-1 showed presence of ER arrays (fig. 5 K1). Interestingly, the aged mice of both strains showed presence of several Golgi apparatus units (fig. 5I ‘Go’). In the MPTP-administered aged C57BL/6J they appeared circular and possessed bloated saccules with “pearl necklace like appearance” (fig. 5, F&H; J&L1; ‘Go’). In the older C57BL/6J apoptotic bodies were noted (fig. 5, I1; ‘Ab’) while in CD-1, the neuronal nucleus was at times crenellated. The neurons of MPTP-injected young CD-1 harbored several lysosomes, which were conspicuously absent in the neurons of young C57BL/6J (fig. 5 B&D ‘Ly’).

#### B: Astrocytes

The astrocytic nuclei were larger but not uniformly ovoid/round like those of oligodendrocytes. Their nuclear chromatin was fine and granular. The cytoplasm was sparse and granular within the perinuclear zone (Luse SA, 1956), but more electron dense than the neuronal cytoplasm. The nucleus remained euchromatic through aging and in response to MPTP. The astrocytes of young and middle aged C57BL/6J had numerous long and tubular cytoplasmic mitochondria (fig. 6, ‘M’, compare A&B and E&F) whereas those in the myelinated axons were spherical (‘Ma’). In CD-1, the cytoplasmic mitochondria were oval (fig. 6, compare C&D and G&H). The astrocytic mitochondria were relatively longer in the old CD-1 (fig. 6, compare E&G). The ER was relatively well maintained through aging and in response to MPTP in both strains. Golgi saccules were semicircular and dilated in the middle aged MPTP-injected CD-1 (fig. 6, Go, compare G&H and K&L). The scale bar is 1μm for all micrographs except ‘K’.

**Figure 6:**
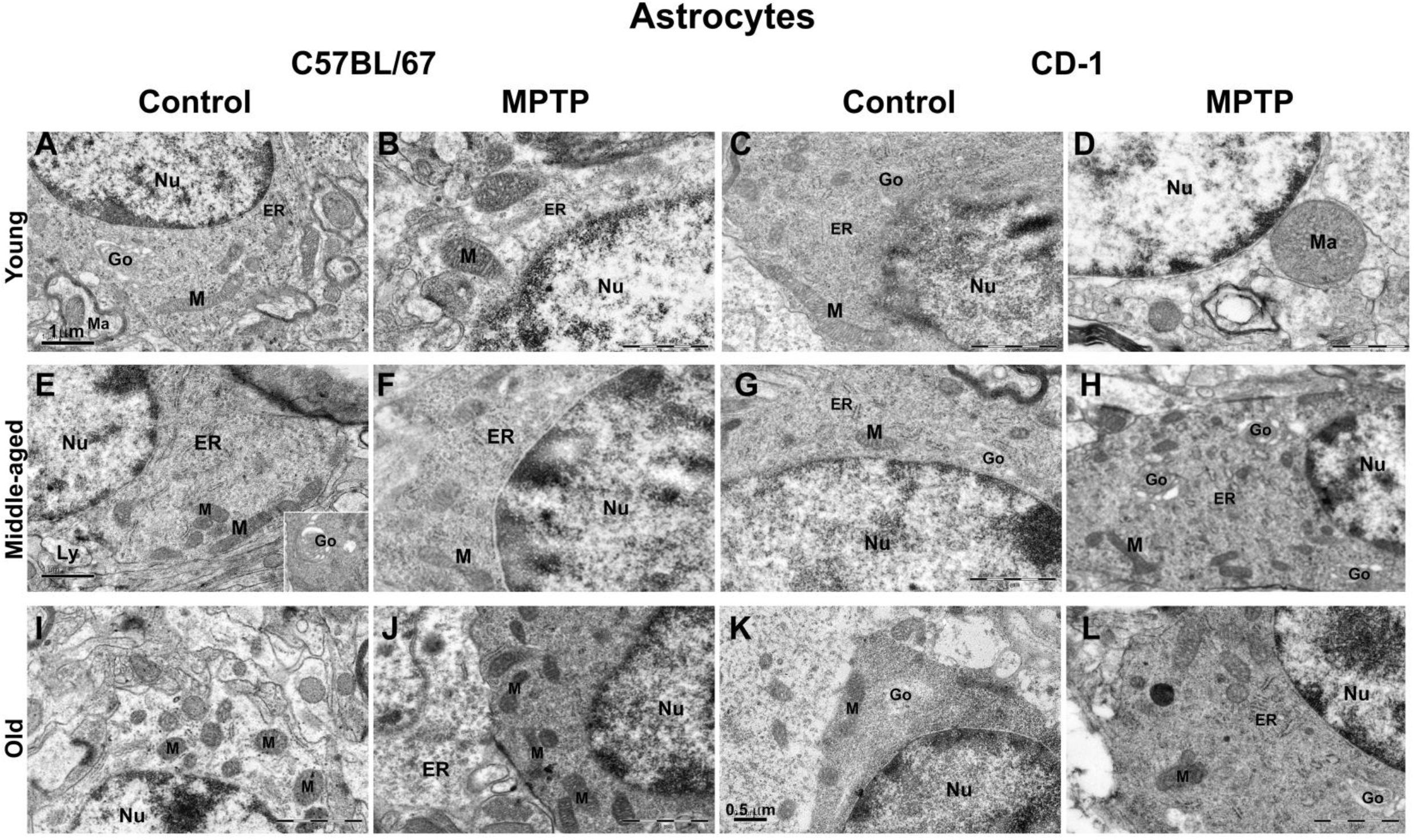
Astrocytic organelles mostly resist age and MPTP induced alterations. Representative electron micrographs of astrocytes at different ages and in response to MPTP.Astrocytes had large and euchromatic nucleus. Note the numerous long and tubular cytoplasmic mitochondria in young and middle aged C57BL/6J (‘M’, A vs. E) and the alterations following MPTP (B vs. F) and spherical ones (‘Ma’) in the myelinated axons; oval mitochondria in CD-1 cytoplasm (control C vs. G; post-MPTP D vs. H). In old C57BL/6J, the mitochondria are smaller than old CD-1 (compare I vs. K). MPTP caused qualitative increase in mitochondrial numbers and their elongation (compare I vs. J). Golgi saccules were semicircular and dilated in the middle-aged CD-1 injected with MPTP (Go, compare G vs. H and K vs. L). The scale bar is 1μm for all micrographs except ‘K’.

#### C: Microglia

Most of the microglia were present near the blood vessels and had electron dense cytoplasm with a bean shaped nucleus. Heterochromatin nets and electron dense pockets were noted along the nuclear perimeter. In the young, the nuclei were euchromatic (fig. 7, A-D; ‘M-nu’) while those of middle aged and old, were electron dense (fig. 7, E-L; ‘M-nu’). In the cytoplasm and neighboring tracks of MPTP-injected young C57BL/6J, long tubular mitochondria were noted (fig. 7B, ‘M’). The young CD-1 showed phagocytotic microglia along with the engulfed cells (fig. 7 D; ‘EC’) as also the MPTP-injected old C57BL/6J (fig. 7 J). Interestingly amoeboid “dark cells” were seen around blood vessels in both strains after middle age (fig. 7E, G and I; ‘DC’) and had thin rim of cytoplasm containing few organelles.

**Figure 7:**
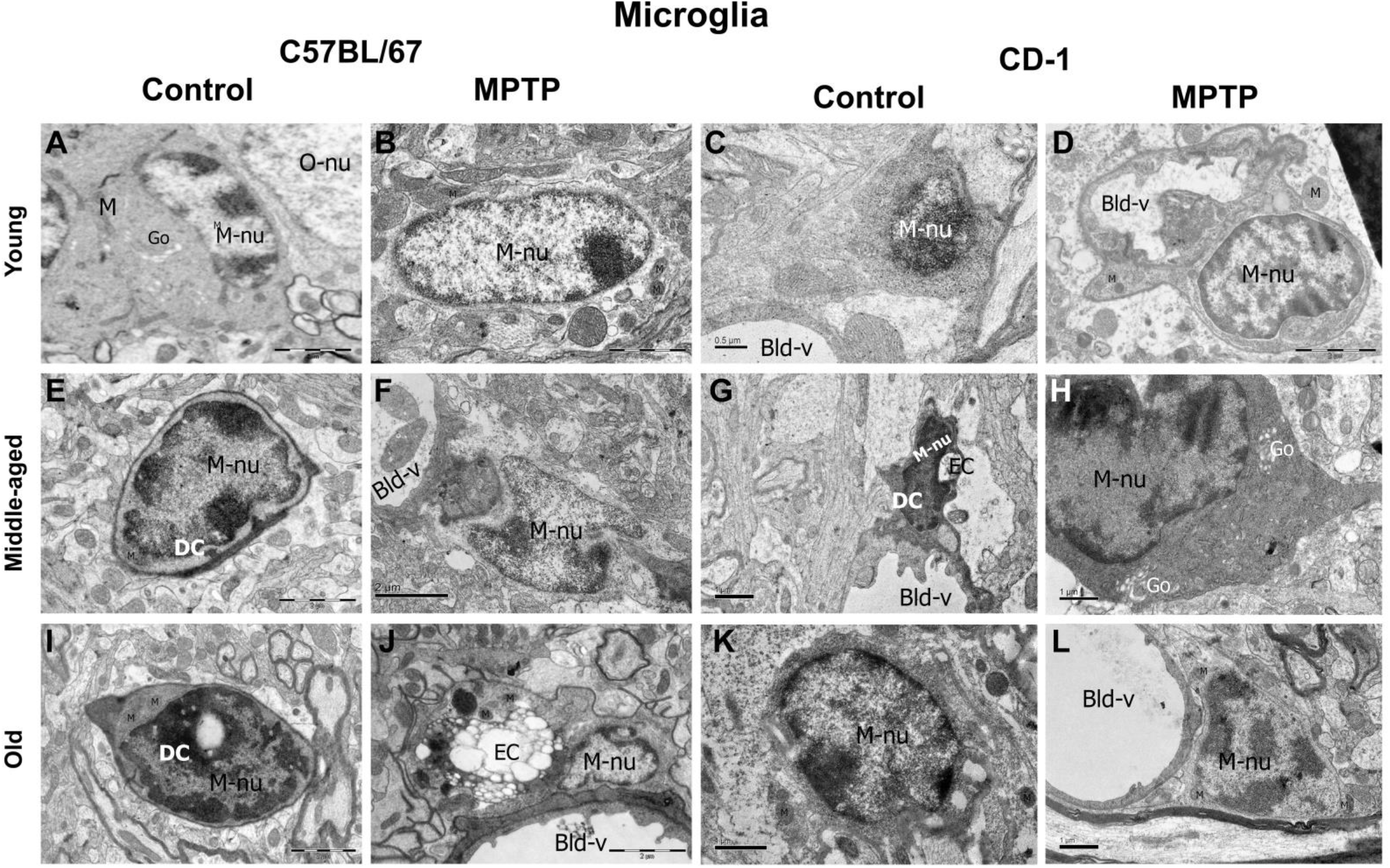
Microglia and Dark cells. Representative electron micrographs of microglia at different ages and in response to MPTP. Most microglia are seen near the blood vessels, have bean-shaped nucleus, electron dense cytoplasm and electron dense pockets along the nuclear perimeter. Note the euchromatic nuclei in young (A-D; ‘M-nu’) and electron dense ones in middle aged and old animals (E-L; ‘M-nu’). In the cytoplasm and neighboring tracks of MPTP-injected young C57BL/6J, mitochondria are long and tubular (B,‘M’). Phagocytotic microglia along with the engulfed cells are seen in young CD-1 (D, ‘EC’) and MPTP-injected old C57BL/6J (J). Amoeboid “dark cells” are seen around the blood vessels in both strains (E, G and I; DC), they have thin rim of cytoplasm containing fewer organelles. The scale bars are specified for each micrograph.

##### Thickening of the vascular basement membrane

Blood vessels showed comparable membrane architecture in the young mice of both strains in control conditions as well as upon MPTP challenge (fig. 8 A-D). At middle age, MPTP injected C57BL/6J showed a membrane discontinuity suggesting a possible breach (E vs F, arrowheads) and thickening of basement membrane in C57BL/6J (arrows, E vs F) but not in CD-1 (F vs H). At old age, MPTP caused thickening of vascular basement membranes in both strains (arrows, I vs J; K vs L). Macrophage-like-cells (** I & J) were seen in the lumen in old C57BL/6J. Dark cells (DC) were noted too.

**Figure 8:**
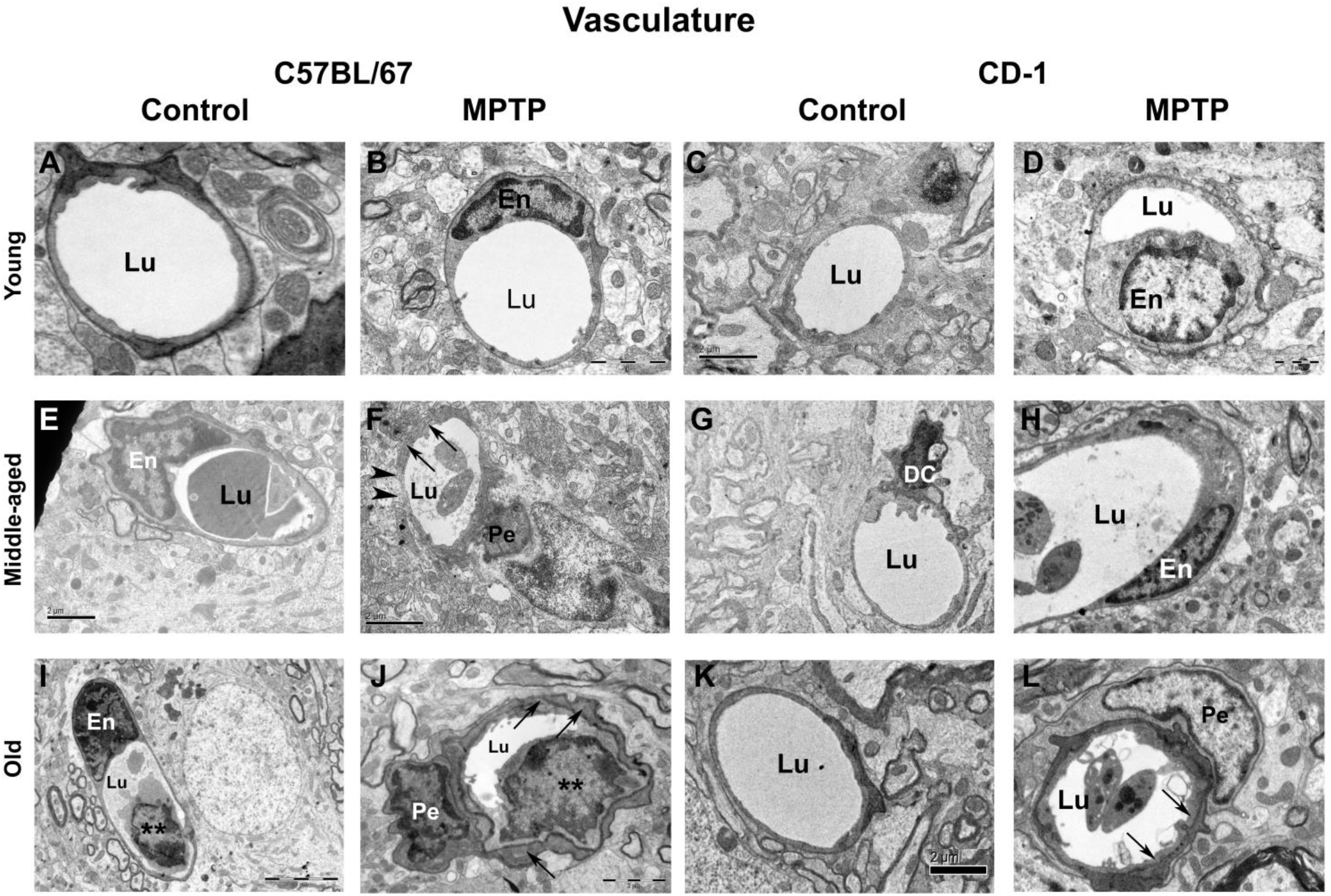
MPTP affects vasculature at middle age in C57BL/6J. Representative electron micrographs of blood vessels from different age groups and in response to MPTP. Note the thin basement membrane of the blood vessels inthe young (A-D). MPTP injected middle-aged C57BL/6J showa discontinuous membrane suggesting a possible breach, (‘F’, arrowheads) and thickening of basement membrane in C57BL/6J (arrows, E vs F) but not in CD-1 (F vs H). Note that at old age, in response to MPTP, the basement membranes of the vessels appear thickerin both strains (arrows, I vs J; K vs L).

###### MPTP caused microbial dysbiosis

## DISCUSSION

### The baseline number of microglia and astrocytes show minor inter-strain differences

Microglia are selectively more populous in the hippocampus and SNpc (Lawson et al., 1990), the target regions of the common age-associated diseases like Alzheimer’s and Parkinson’s disease respectively; endorsing their role in neurodegeneration and susceptibility. In aging mice, microglial priming is equated to an immune vigilant state that results in their de-ramification, hypertrophy, increase in numbers and exaggerated response to sub-threshold challenges. They synthesize ROS and pro-inflammatory cytokines like TNF-α, IL-6, IL-1β; while curtailing the synthesis of anti-inflammatory cytokines (Damani et al., 2011; Grabert et al., 2016). The presence of more microglia at middle age corroborates with the possibility of a pre-inflammed status and presence of damage associated molecular patterns (DAMPs) in-situ, thereby inflammasomes too may be vital in determining susceptibility. We found that normal C57BL/6J mice have fewer substantia nigra neurons (Vidyadhara et al., 2017) and more baseline apoptosis (Yarreiphang et al., unpublished data) vis-à-vis the CD-1 mice. Neuronal apoptosis has been shown to flag the entry of microglial precursor cells into the zebra fish brain (Casano et al., 2016). Thus, just moderately more microglia (8-18%) in the control C57BL/6J substantia nigra suggests a partial influence of numbers on susceptibility.

Aging selectively affects the already sparse astrocytes in the substantia nigra, as against the other mesencephalic niche (Damier et al., 1996). Under physiological conditions, they secrete trophic factor GDNF (Grondin et al., 2003), therefore the presence of moderately fewer astrocytes in C57BL/6J (10-15% less than CD-1) and an acute increase in their numbers at middle-age that persisted till old age, suggest age-related reduction in neuroprotection and gliosis-assisted inflamm-aging. The latter could be linked to cytokines. Whereas, the sedate increase in astrocytic numbers in CD-1 implicate graded changes in the milieu.

### Higher TNF-α expression in the susceptible strain may be a steering factor

DA neurons are sensitive to the pro-inflammatory cytokine TNF–α (Sriram et al., 2006); which is primarily secreted by microglia and in synchrony with IFN-γ/IL-1β induces neurodegeneration (Chao et al., 1995; Jeohn et al., 1998). Thereby, the dying neurons hold microglia in their cytotoxic state, to escalate synthesis of TNF-α leading to a self-perpetuating neuroinflammation. The higher levels of TNF-α released by the glia recruit their neighbors, to set off a vicious cycle, wherein the pro-inflammatory environment becomes self-propagating (Noh et al., 2014). Although only mildly higher in numbers, the persistently higher baseline TNF-α level through aging in C57BL/6J, suggest that in their nigra, the microglia may be consistently primed, contributing to neuronal susceptibility. Elevated TNF-α levels are reported in mouse models as well as in autopsied brain tissue/CSF of PD patients (Mogi et al., 1994). Low levels of TNF-α are protective (Chertoff et al., 2011). Thus, the lower levels in CD-1 in addition to being a mark of low susceptibility may be potentially neuroprotective. Our auxiliary finding of TNF-α up-regulation in middle-aged C57BL/6J, indicates that an imbalance in cytokine milieu precedes senescence.

IL-6, a pleiotropic cytokine, is up-regulated in SN, CSF and serum of PD patients (Blum-Degena et al., 1995; Hofmann et al., 2009). Conversely, Bolin et al., (2002) reported amplified MPTP-susceptibility in IL-6^-/-^ mice. In our study, since IL-6 followed a similar age-related expression pattern in both strains, it may not dictate baseline susceptibility. Yet, the differences following MPTP validate its role as a toxicity signal. The middle-age demarcates a period of enhanced susceptibility. MPTP-induced IL-6 expression in old C57BL/6J at later stages of the challenge, suggests a link with astroglial responses or excessive neuronal death.

Reactive microglia in the degenerating SN overproduce IL-1β, to activate astrocytes and promote iNOS secretion (Chhor et al., 2013). Chronic IL-1β expression in Wistar-rat substantia nigra induced progressive neurodegeneration, microgliosis and motor disabilities (Ferrari et al., 2006). The acute increase in IL-1β at d1 post-MPTP that persists till d4 in the young C57BL/6J vis-à-vis CD-1 endorses immediate microglial priming and activation in the former. Thus, the mice differ in the immediacy or respondence indices. Moreover, noticeably higher basal TNF-α in C57BL/6J at middle and old age underlines its role in basal susceptibility, aging and neurodegeneration, whereas, IL-6 and IL-1β appear to be weak responders to MPTP.

### The resistant strain has higher TGF-β through aging, but lower IL-10 levels at middle age

TGF-β1 inhibits microglial activation and protects DA neurons against MPTP-toxicity (Arimoto et al., 2007; Pintado et al., 2011). The low baseline TGF-β expression in C57BL/6J through aging and following MPTP, suggest that this sensitive strain is ill equipped against neuroinflammation. Since CD-1 have more substantia nigra DA neurons and many of them resist MPTP (Vidyadhara et al., 2017), higher TGFβ1 levels allude to neuroprotection and DA neuronal survival through aging.

IL-4 protects DA neurons against MPP^+^ toxicity by up-regulating CD200, a microglial resting signal (Lyons et al., 2009). The moderate upsurge at d4 in MPTP-injected young CD-1, suggests an auxiliary course by the microglia (Hirsch and Hunot, 2009). In view of its pronounced increase in basal levels of the MPTP-sensitive C57BL/6J at middle-age, it is likely that IL-4 may also have a pro-inflammatory role. Bok et al., (2018) while showing the microglia-specific expression of IL-4 demonstrated an LPS-induced up-regulation in expression and rescue of substantia nigra DA neuronal loss by antibodies against IL-4.

IL-10 stimulates CD200 expression in neurons and induces astrocytes to synthesize anti-inflammatory TGF-β. It inhibits microglial synthesis of TNF-α, NO and ROS, in-vitro, to neutralize oxidative stress (Balasingam and Yong, 1996; Ledeboer et al., 2002). The up-regulated baseline IL-10 expression in C57BL/6J and upon MPTP at middle/old-age may be a compensatory increase. However, these increases do not parallel a raise in TGF-β levels; hinting at a failed rescue attempt.

Thus, the noticeable increase in basal levels of both TNF-α and IL-6 as well as the anti-inflammatory IL-1β and IL-10 at middle age; expound a hovering imbalance of pro- and anti-inflammatory cytokines, therefore tempting one to speculate that middle-age imitates the prodromal period in the susceptible strain.

### Strain specific variability in enzyme responses

MAO-B inhibitors are promising candidates in the treatment of early-PD (Rabey et al., 2000). MAO-B level increases with age (Irwin et al., 1997) and is doubled in SN of PD patients (Damier et al., 1996). The age-associated up-regulation in MAO-A and MAO-B levels in C57BL/6J suggests a functional decline. The gradual age-related reduction in CD-1 may be a natural phenomenon that assists in combating MPTP-related stress. Although the reasons for higher basal level of MAO-A and MAO-B in young CD-1 are unclear, it may be due to higher number of neurons or DA terminals or it may be a bystander susceptibility marker. MPTP may cause a feed-forward effect on glia, triggering off a toxicity cycle.

HO-1 a cytoprotective, anti-apoptotic, and anti-inflammatory enzyme; down-regulates pro-inflammatory cytokines like TNF-α1 and IL-1β and up-regulates anti-inflammatory cytokine IL-10 *in-vitro* (Doré et al., 1999; Petrache et al., 2000). Predominantly expressed by astrocytes (Dwyer et al., 1995), it is positively correlated with aging and PD (Schipper et al., 1998). It was projected as a potential biomarker due to high levels in the patient saliva(Song et al., 2018). HO-1 overexpression tendered neuroprotection in MPP^+^ treated Parkinsonian rats via BDNF and GDNF (Hung et al., 2008). Thus, the gradual age-related increase in its levels in C57BL/6J and in response to MPTP may indicate a cellular offset response to oxidative stress. Induction of iNOS causes NO release, which when protracted triggers oxidative damage in DA neurons (Nathan and Xie, 1994). The MPTP-induced up-scaling of iNOS in both strains supports this hypothesis.

The higher baseline iNOS and MAO-B levels alongside lower HO-1 levels in CD-1 are presently un-explained. Alternatively, these may be markers of sub-threshold susceptibility in them.

Neurons secrete the chemokine fractalkine (CX3CL1), to maintain microglia in resting state (Cardona et al., 2006). The inherently higher fractalkine levels in CD-1 indicate their healthy status. Age-related decrease in C57BL/6J relays enhanced phagocytic signals, an indirect indicator of neuronal loss. Overexpression in response to MPTP in young mice implies the activation of compensatory responses during youth which decline with age.

### Surviving neurons harbor strain-specific ultrastructural signatures

This is the first study on the alterations in the ultrastructureof mice substantia nigra with aging and in response to MPTP. The age-related and MPTP-induced shrinkage of mitochondria in C57BL/6J validates the ensuing mitochondrial dysfunction and aging as a risk factor for PD. We earlier found higher DRP-1 levels in the lateral/ventral substantia nigra of C57BL/6J, earmarking the inherent susceptibility of the mitochondria. On similar grounds, upregulation of mitochondrial fission protein dynamin-like protein 1 (DLP1/DRP1) as well as down-regulation of fusion proteins Mfn1 and Mfn2 were noted in the substantia nigra of PD patients (Zhao et al., 2017).

The relatively better-preserved mitochondrial structure with age and upon-MPTP in CD-1 complements the higher HSD-10 (mitochondrial fusion-associated protein) expression (Seshadri and Alladi, 2019). Elongated mitochondria are deft in energy generation (reviewed by Galloway et al., 2010) and calcium uptake (Lewis et al., 2018). In *Caenorhabditis elegans*, modulation of mitochondrial proteases SPG-7 and PPGN-1 enhanced mitofusion (Chaudhari and Kipreos, 2017) to extend their overall survival. Thus enhanced HSD-10 expression (Seshadri and Alladi, 2019) and elongated mitochondria in old CD-1 neurons may be the survival modalities, which may be lacking in C57BL/6J.

Both fragmented and dilated ER are major pathological notations in A53TαS Tg mice substantia nigra(Colla et al., 2012)and rotenone model of sub-cutaneous administration(Zhang et al., 2017), suggesting ER dysfunction in PD. The MPTP-induced ER shortening in C57BL/6J and its dilation in CD-1 suggests existence of strain-specific differences as well as different aspects of ER dysfunction in PD, which needs to be studied in detail. The presence of ER arrays in old CD-1 could be rejoinders of enhanced protein synthesis to compensate for the concurrent protein loss or mis-folding.

The presence of many intact Golgi units in the aged substantia nigra of both strains suggest a compensatory increase to circumvent age effects on protein packaging and post-translational processing. The “pearl necklace like globose” bloated saccules of Golgi units upon MPTP-injection in old C57BL/6J, hint at disease-induced functional impairment. Knockdown of adhesion proteins like GRASP 55/65 cause Golgi cisternae swelling(Lee et al., 2014). Simulation studies suggest that aberrations in biophysical properties like osmotic pressure, adsorption and adhesion energies, and precise vesicle addition frequency in addition to biological properties like rim stabilizer proteins cause abnormal self-organization into circular/fused Golgi complexes (Tachikawa and Mochizuki, 2017). The presence of lysosomes in neurons of MPTP-injected young CD-1 that were visibly fewer in C57BL/6J may suggest either the activation of lysosomal/UPR pathway or lysosomal accumulation due to impaired late endocytic pathway (Guerra et al., 2019). The apoptotic bodies in older C57BL/6J, signal the occurrence of age-associated apoptosis. The crenellated nuclei in MPTP-injected old CD-1, suggest necroptosis as seen in striatal cells of a Huntington’s mouse (Turmaine et al., 2000). Thus, a majority of organelles were affected in C57BL/6J both with aging and upon MPTP indicating affliction of associated cellular processes. Contrarily, elongation of mitochondria, preservation of Golgi apparatus and presence of lysosomes may be pointers of resilience in CD-1. Amongst the organellar defects, ER dilation is a sure sign of susceptibility in CD-1.

### Elongation of glial mitochondria, a distinctive feature of pathogenesis

In an interesting spin off, unlike the neurons, ultrastructure of glial organelles was preserved during aging and following MPTP. The glial mitochondria that were smaller and fewer in controls C57BL/6J, appeared enlarged/elongated in response to MPTP. Hoekstra et al., (2015) showed a reduction in fission protein DLP1/DRP1 in both neurons and astrocytes in PD cortex along with fused elongated mitochondria in primary cortical astrocytes transfected with DLP1-siRNA. Co-culturing them with cortical neurons caused neuronal atrophy and excessive calcium release; suggesting neurotoxic effect of fused astrocytic mitochondria. In a stroke model, the penumbral astrocytes displayed hypertrophic and polarized processes; elongated mitochondria and lost their neuroprotective ability (Fiebig et al., 2019). Overexpression of mutant ubiquitin (UBB+1) protected astrocytes from oxidative stress and H_2_O_2_-induced cell death by destabilizing mitochondrial fission-specific proteins, leading to mitochondrial fusion (Yim et al., 2014). Although the exact corollaries are not clear, the combination of astrogliosis and increased pro-inflammatory cytokines; tempts one to speculate that elongated mitochondria assist astroglial propagation into neurodegenerativesequels.

Mitochondrial elongation in microglia of MPTP-injected young C57BL/6J mice, may too have neurodegenerative consequences. LPS-activated mouse cerebral microglia, stimulate DRP-1 and ROS synthesisin-vitro, to elongate tract borne mitochondria (Katoh et al., 2017). The dilated Golgi apparatus in microglia indicate that while aggravating neuroinflammation, the microglial protein packaging process is also affected. Under chronic stress, in aging and in AD; microglia had condensed, electron dense cytoplasm and nucleoplasm which imparted a striking “dark” appearance (Bisht et al., 2016). Dark cells noted in both strains from middle age, may have similar pathological objective. In most cells, the cytoplasm appeared as a thin rim, reducing the scope to visualize other organelles.

Our findings of mitochondrial elongation may have clinical implications. For instance, the antioxidant coenzyme Q10 (CoQ10) supports mitochondrial function while reducing the DA neuron loss in an animal model of PD (Spindler et al., 2009), yet it failed in the phase III clinical trials (The Parkinson Study Group QE3 Investigators 2014). Similarly, CoQ10 derivative MitoQ also failed (Snow et al., 2010). It is likely that these molecules stabilized the glial mitochondria too, thereby nullifying the neuronal outcome. It is vital to study the glial mitochondrial responses in isolation and on a temporal scale, to better understand the phenomenon.

### Earlier age at onset of MPTP-induced basement membrane features in C57BL/6J

MPTP caused basement membrane thickening similar to that reported in PD (Farkas et al., 2000). Presence of gaps in the middle aged C57BL/6J and old CD-1 implies that the blood-brain barrier (BBB) is vulnerable earlier in life in C57BL/6J, an additional signal of negative impact of aging or on the gut microbiota (Montagne et al., 2015) or of inherent susceptibility.

## CONCLUSION

In summary, neuro-glial interactions, senescence related changes in glia, cytokine levels etc. are vital determinants of neuronal survival and differ greatly in the MPTP-resistant C57BL/6J and MPTP-susceptible CD-1 white mice strain. The intended use of male animals for the study may be a limiting factor, however in view of the male preponderance of the disease; our observations provide vital clues of disease pathogenesis. Besides, female mice are also known to show higher fatality in response to MPTP, due to differences in peripheral metabolism of MPTP; independent of its effects on dopaminergic neurons. It is also pertinent to compare these factors between male and female animals to understand the premise for neuroprotection in females.

Our findings in general, may be extrapolated to different human populations that are either vulnerable or resistant to PD. For instance, in our study, the CD-1 represents Asian-Indians who have inherently lower prevalence rate than the Caucasians. The prominent differences in pro- and anti-inflammatory cytokine levels at middle-age suggests that by design, the middle-age milieu is acquiescent to neurodegeneration and may well be the critical soft period for the onset of neurodegenerative diseases. It is likely that senescence may result from differences in the in-situ inflammasomes, which merit detailed investigations. The pathogenesis is effectively assisted by the diabolic differences between the neuronal and glial mitochondria. Thus, glia are major players in aging and disease and may explain the ethnic bias in prevalence of PD.

## Abbreviations

MPTP: 1-methyl-4-phenyl-1, 2, 3, 6-tetrahydropyridine
BBB: Blood brain barrier
BSA: Bovine serum albumin
CoQ10: Coenzyme Q10
D1: day 1 post-MPTP:
D4: day 4 post-MPTP
D7: day 7 post-MPTP
DAMP: Damage associated molecular patterns
DLP1/DRP1: Dynamin-like protein 1
ELISA: Enzyme-linked immunosorbent assay
ER: Endoplasmic reticulum
GFAP: Glial Fibrillary Acidic Protein
H_2_O_2_: Hydrogen peroxide
HO-1: Hemeoxygenase-1
Iba-1: Ionized calcium-binding adaptor protein-1
IFN-γ: Interferon-γ
IL-1β: Interleukin-1β
iNOS: Inducible nitric oxide synthase
Ir: immunoreactive
MAO-A: Monoamine oxidase A
MAO-B: Monoamine oxidase B
MPP^+^: 1-methyl-4-phenylpyridinium
PD: Parkinson’s disease
PVDF: Polyvinylidene difluoride
RT: Room temperature
SNpc: Substantia nigra pars compacta
TMB: 3,3’,5,5’ -tetramethylbenzidine
TNF-α: Tumor necrosis factor-α

## Declarations

### • Ethics approval

All the experimental protocols on mice were approved by the Institutional Biosafety committee and Institutional (NIMHANS) Animal Ethics Committee, in accordance with the guidelines of the CPCSEA, India, and NIH, USA.

### • Consent for publication

Not applicable

### • Availability of data and materials

The raw datasets used and/or analysed during the current study are available from the corresponding author on reasonable request.

### • Authors’ contributions

APL contributed to study design, performed experiments and collected data and provided 1^st^ draft; UB performed experiments and collected data; MP performed statistical analysis of the data; RSK & BKCS performed electron microscopy and data analysis; TRR and BMK: contributed to data analysis and revision of MS; PAA conceptualized the study, performed data analysis, edited the manuscript and obtained funds.

## • Acknowledgements

We thank Dr. M.M. Srinivas Bharath, Head, Department of Clinical Psychopharmacology and Neurotoxicology; Dr. Gayathri N, Department of Neuropathology for laboratory facilities.

